# Migratory herbivorous waterfowl track multiple resource waves during spring migration

**DOI:** 10.1101/2023.04.14.536842

**Authors:** Fei Xu, Wei Wu, Jie Wei, Qinchuan Xin, Ben Wielstra, Frank A. La Sorte, Zhijun Ma, Guangchun Lei, Jialin Lei, Wenzhao Wu, Yongchuan Yang, Peng Gong, Bing Xu, Yali Si

## Abstract

East Asian herbivorous waterfowl intensively use farmland in spring, next to their natural habitat. Accordingly, they might have expanded their migration strategy from merely tracking the green wave of newly emerging vegetation to also incorporating the availability of post-harvest agricultural seeds (here dubbed the seed wave). However, if and how waterfowl use multiple food resources to time their seasonal migration is still unknown. We test this migration strategy using 167 spring migration tracks of five East Asian herbivorous waterfowl species and mixed-effect resource selection function models. We find all study species arrive at their core stopover sites in the Northeast China Plain after agricultural seeds become available, extend their stay after spring vegetation emerges, and arrive at breeding sites around the emergence of vegetation. Moreover, the cue used in exploiting the availability of both food resources varies in different regions and among species. At the core stopover sites, larger-bodied species use the seed wave and smaller-bodied ones adopt the green wave. At the breeding sites, the former follow the green wave, and the latter track the snow-free time. Our findings suggest that waterfowl track multiple resource waves to finetune their migration, highlighting new opportunities for conservation.

## Introduction

Migratory species move between their breeding and non-breeding grounds and benefit from the exploitation of seasonal food resources across the annual cycle [1]. Foraging opportunities occur in a series of pulses, at different times and places, which generate “resource waves” across the landscape [1–5]. The green wave hypothesis posits that the migration progress of herbivores is driven by the seasonal growth of new vegetation at stopover sites along the migration route [6, 7]. Studies have verified this hypothesis for a variety of herbivorous animal groups, such as geese [4, 6, 8-10] and ungulates [11–13]. Not only individual-level information compiled using tracking technology [6] but also population-level information compiled by large-scale community science initiatives [14] support such hypothesis. During vernal migration, birds mainly arrive at their stopover sites after the emergence of spring vegetation, but can reach their breeding grounds earlier, thus overtaking the green wave, and allowing their offspring to benefit from the peaks of food resources [4, 9]. While herbivorous species have been documented to track the green wave in Northern Europe [6, 10, 15] and North America [14], this has not been the case for many herbivorous waterfowl populations worldwide, especially within East Asia [15].

Due to extensive land cover change worldwide, herbivorous waterfowl have expanded from using natural ecosystems to exploiting food resources in agricultural landscapes [16, 17]. For example, during the spring-staging period, herbivorous waterfowl have been observed to consume new sown cereals in Europe [18], post-harvest corn remaining in cornfields, and spilled grain in rice paddies in North America [19–21] and in Asia [22–27]. This way they benefit from the post-harvest management of agricultural lands [28, 29]. This is particularly the case for East Asian waterfowl populations, where in China large areas of natural wetland have been converted into farmland since the 1950s [30–33]. As a consequence, East Asian waterfowl might take the availability of both food resources into account during spring migration. However, this possibility of tracking multiple resource waves during migration has not yet been investigated.

Migratory herbivorous waterfowl use the natural habitat (e.g., wetlands, grasslands, shrublands) and farmland habitat differently. Since many waterbodies and grasslands in East Asia are semi-natural, we use the term ‘natural habitat’ hereafter to represent areas where birds feed on spring sprouts of vegetation (representing the green wave). Within farmlands, East Asian waterfowl primarily consume seeds, rather than crop vegetation, e.g., spilled grains and leftover maize that remained after the harvest [22] in the previous fall and so have been carried over into spring. These seeds have been found to offer a higher food quality compared to natural vegetation [16]. Spilled seeds are only available after the harvest and during the period that the ground is not frozen, resulting in two waves in autumn and in spring, that we here dub the seed wave (Figure S1). In winter at the core stopover area Northeast China Plain, the daily minimum temperature can drop below −25 ℃ [34, 35]. It often starts snowing from November and ends around the end of March [34, 36]. This makes the post-harvest seeds not accessible in winter but these seeds can be used by waterfowl after the ground thaws around the end of March [36]. The seed wave then ends at the onset of sowing in early May [36], when the new seeds that are sowed covered with pesticides (Figure S2*C*). Thus, the seed wave in spring in the core stopover area starts around the end of March (with the end of snowmelt as a surrogate) and ends in early May with the start of sowing (Figures S1 and S2) [37]. The timing of seed availability starts before, and partially overlaps with, the emergence of vegetation [15]. How would the herbivorous waterfowl time their migration based on these two resource waves is still unclear.

We aim to investigate if and how herbivorous waterfowl track multiple resource waves during spring migration. We analyze 167 spring migration tracks from 2015 to 2021, derived from satellite tracking of 99 individuals for five herbivorous waterfowl species (swan goose *Anser cygnoides*, tundra bean goose *A. serrirostris*, greater white-fronted goose *A. albifrons*, lesser white-fronted goose *A. erythropus*, and tundra swan *Cygnus columbianus*) in East Asia, using a mixed-effect resource selection function model. We hypothesize that migratory herbivorous waterfowl select resources upon arrival at stopover and breeding sites based on the timing of food resource availability in both natural habitat and farmland (Figure 1). We predict that: 1) All study species time their migration to use both resources (seeds and natural vegetation) at their core stopover sites. They use farmland more frequently upon arrival at the core stopover sites, and shift to natural habitat after the vegetation emerges. All study species arrive at their natural habitat in the breeding sites around the time of vegetation emergence; 2) Waterfowl tend to track the start of seed availability (i.e., the seed wave) at their core stopover sites and then follow the newly emerged plants (i.e., the green wave) as a cue to arrive at their breeding sites (Figure 1). Understanding how herbivorous waterfowl use the spatiotemporal availability of multiple food resources during migration is fundamental to predict species’ responses to land cover and climate change. The wider implications of our findings are new opportunities for effective management of multiple types of habitats to conserve migratory waterfowl.

**Figure 1.**
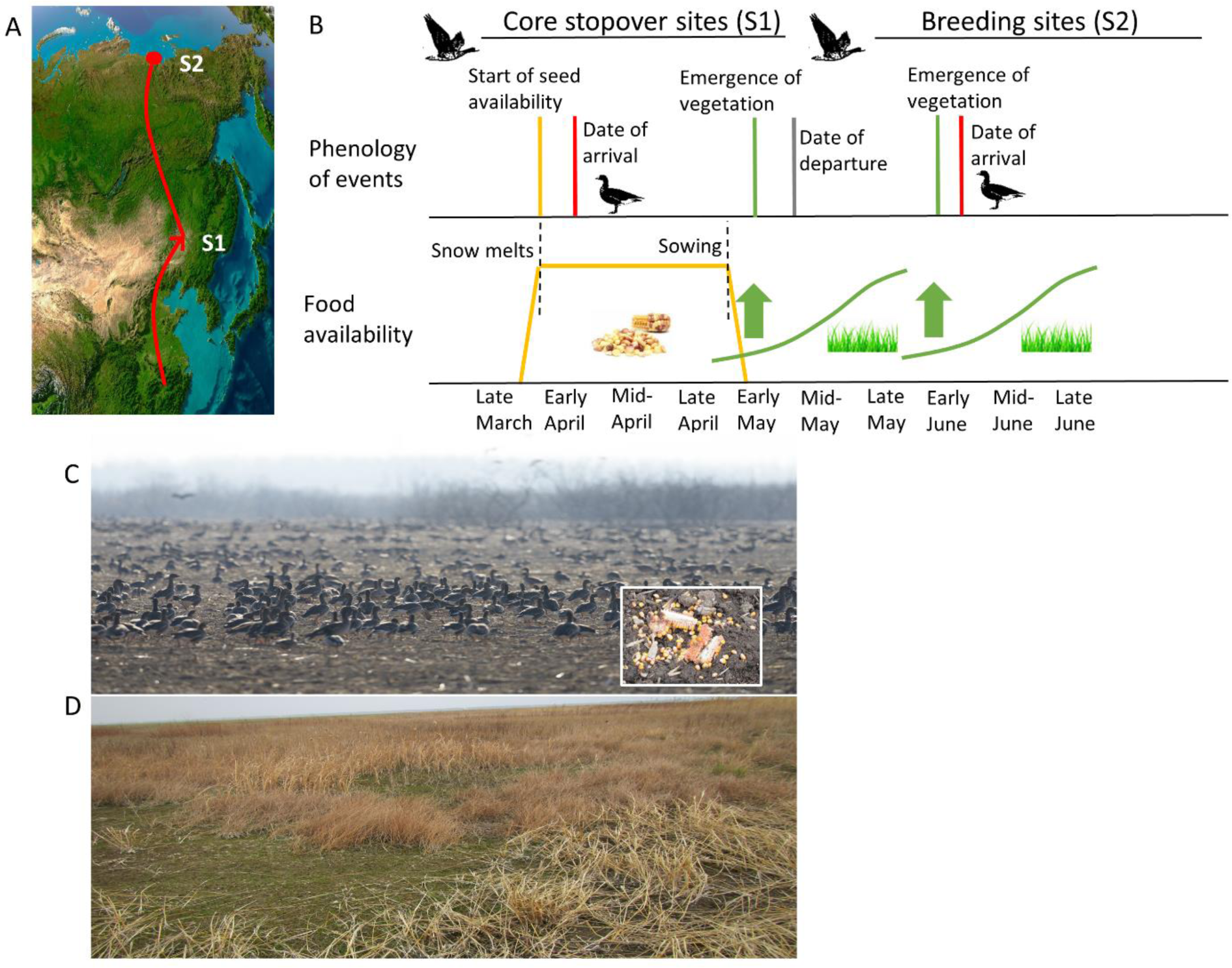
Schematic diagram of tracking multiple resource waves by East Asian migratory herbivorous waterfowl. **(A)** Core stopover sites (S1) and breeding sites (S2) along the spring migration route. **(B)** During spring migration, birds arrive on farmlands at the core stopover sites after the start of seed availability (seed wave; orange line), extend their stay after spring vegetation in natural habitats emerges (green wave; green lines) (S1) and reach the breeding grounds after the emergence of vegetation (green wave; green lines) (S2). **(C)** Photo of greater white-fronted geese (*Anser albifrons*) and tundra bean geese (*A. serrirostris*) foraging in farmlands (taken 19 March 2021) on Yuquan island, Hunchun, Yanbian Korean Nationality Autonomous Prefecture, Jilin, China (photo credit: Haixiang Zhou). Inset photo shows leftover maize (taken 2 May 2016) on farmlands in Fuyu Village, Nehe, Qiqihar, China (photo credit: Jie Wei). **(D)** Photo of new spring vegetation (taken 1 May 2016) embedded in old vegetation in Baishan town, Longjiang, Qiqihar, China (photo credit: Jie Wei).

## Materials and Methods

### Analysing bird migration patterns

We analyzed the satellite tracking data (i.e., 167 spring migration tracks) of five East Asian herbivorous waterfowl species from 2015 to 2021, covering 22 tracks of swan geese from 19 individuals, 8 of tundra swans from 4 individuals, 54 of tundra bean geese from 28 individuals, 63 of greater white-fronted geese from 34 individuals, and 20 of lesser white-fronted geese from 14 individuals. Lesser white-fronted geese (*Anser erythropus*), greater white-fronted geese (*A. albifrons*), tundra bean geese (*A. serrirostris*), and tundra swans (*Cygnus columbianus*) were captured and tagged at their East Dongting Lake (29°N, 113°E) and Poyang Lake (29°N, 116°E) wintering sites in the Yangtze River Floodplain, China during the 2014 to 2018 wintering seasons. Swan geese (*A. cygnoides*) were captured at the breeding site Hulun Lake, Hulun Buir, China (48.3°N, 117.4°E) during the 2017 molting season. All birds were equipped with GPS - GSM (Global Positioning System – Global System for Mobile Communications), solar-powered loggers. GPS locations were recorded every 2 h. Information used in the analysis on migration patterns include Bird ID, track year, longitude, latitude, and time of the GPS location. Detailed methods on birds capture and deployment of the satellite transmitters can be found in Si, Xu [22] and Lei, Jia [38].

We quantified the migration parameters (Table S1) in three steps. First, we subset all GPS locations between January 1st to July 31st, thereby including months of the whole spring migration period (February-June) and the months in which all five herbivorous waterfowl have arrived at breeding sites (June-July). The definition of the spring migration period is that GPS locations recorded from the day that birds left their wintering sites to the day that they reached their breeding sites [8]. Second, we labeled daytime and nighttime locations, by identifying the sunrise and sunset time for each location based on algorithms provided by the National Oceanic & Atmospheric Administration (NOAA) (https://www.esrl.noaa.gov/) [39]. Third, we identified stopover sites and the length of stay in each site through the space-time permutation model (Supplementary Text) in the SaTScan statistics (http://www.satscan.org) [22, 40]. We defined the stopover sites as those sites where birds stayed for at least two days [41] and moved within a radius of less than 50 km [22, 41-43]. The arrival/departure time of the wintering/stopover/breeding sites for each individual in each track year were defined as the first day it arrived/left a specific site in which it showed a continuous presence/absence [23]. Cumulative length of stay in the Northeast China Plain and Russian stopover sites was defined as the sum of all days spent at stopover sites by an individual in its respective region. Migration distance was defined as the cumulative distance travelled between the wintering site and breeding site [22].

We calculated mean dates of migration timing upon arrival at or departure from wintering sites, core stopover sites, and breeding sites for all tracks of each species, i.e., 1) mean departure date from the wintering grounds, 2) mean arrival date at core stopover sites, 3) mean departure date from core stopover sites, and 4) mean arrival date at breeding sites. For each of the four types of migration timing, we compared the dates among five herbivorous waterfowl species in relation to their body size, which was obtained from EltonTraits 1.0 [44]. Differences on the migration timing across species were assessed using one-way ANOVA followed by Tukey’s honestly significant difference (HSD) multiple comparison.

### Investigating resource utilization in migration

To examine the habitat use of tracked birds along the spring migration route, we used two land cover products. The 30 m spatial resolution spring land cover product [45] from 2015 was selected to capture the natural habitats and farmlands within China. The European Space Agency Climate Change Initiative (ESA-CCI) 300 m spatial resolution land cover product from 2015 (http://maps.elie.ucl.ac.be/CCI/viewer/index.php) was used to capture the relatively homogeneous landscape within Russia. Two classification systems from the two products were aggregated into seven land cover types: grassland, farmland, water, wetland, forest, shrubland, and ‘other’ (i.e., barren land, lichens and mosses, snow and ice, or impervious). Habitat use at each stopover site was estimated using the percentage of bird GPS locations on each land cover type [22]. Habitat use at each latitude was estimated using the percentage of bird GPS locations on each land cover type across all stopover sites (their rounded centroid coordinates) located in the same latitude. The proportion of land cover types in use at each latitude was then calculated as the percentage of bird GPS locations on each land cover type divided by the sum of the percentage of bird GPS locations across all land cover types at the corresponding latitude.

As waterfowl mostly forage during the day, we also calculated the habitat use at daytime (i.e., the proportion of land cover in use during the day) across the whole spring migration period, to better interpret how waterfowl use the food resources across time. To better investigate the multiple resource use across time within the core stopover sites in the Northeast China Plain, we divide the length of stay (Table S1 and Figure S3) of herbivorous waterfowl into two periods, i.e., 1) from the arrival date at each stopover site in the Northeast China Plain to the local emergence of vegetation, and 2) from the local emergence of vegetation till they departure from this region. We then calculate the foraging distribution (i.e., the proportion of GPS locations during the day on farmlands and natural habitats) for all study species during these two different time windows (Figure 3 and Figure S7).

### Calculating the timing of the seed wave and the green wave

For each track, the spring vegetation emergence time in natural habitats and the start of seed availability time on farmlands were calculated. We used the second-order derivative [46, 47] of the smoothed time series of Enhanced Vegetation Index (EVI) derived from the Moderate Resolution Imaging Spectroradiometer (MODIS) Terra 8-day 250 m spatial resolution surface reflectance product MOD09Q1 to extract the spring vegetation emergence time (details are provided in Supplementary Text). Only pure pixels [48] where the proportion of herbaceous area (i.e., wetlands, grasslands, and shrublands) was larger than 0.5 within a 250 m grid cell were used (details are provided in Supplementary Text and Figure S9).

The start of the seed availability time was determined by the end of snowmelt in spring because after the snow melt seeds (remaining from previous year’s harvest) could be easily dug out from the ground. Following the method of Rickbeil, Coops [49] we used the MODIS Terra Snow Cover daily 500 m spatial resolution product MOD10A1 to measure the different degree of snowmelt (i.e., the onset and the end of snowmelt). We first created a time-series to capture the daily percent of snow cover (from day 1 to day 210) for each stopover and breeding site, for each track. We then applied three-piece segmented regression to extract the onset and the end of snowmelt for each pixel (Figure S10). To facilitate interpreting the resource use, in addition to the vegetation availability date and the seed availability date, we also calculated the onset of snowmelt and the end of snowmelt at each site, for each individual bird, in each track year.

### Determining the resource availability – bird arrival relationship

We used a resource selection function (RSF) framework [50, 51] to investigate to what extent the start of resource availability can be matched with the arrival date at the stopover or breeding sites for migratory herbivorous waterfowl during spring migration. Using a logistic regression of presence/absence points, RSFs can be used to define the selection of resources on the day when waterfowl arrive at a specific site [23]. The presence points were defined as all GPS locations recorded on the day of arrival. The absence points were drawn randomly on the date before and after arrival at a specific site during the spring migration period. For each pixel in the specific site, we calculated the number of days between each of the four indices (i.e., the spring vegetation emergence, the start of seed availability, the onset of snowmelt, and the end of snowmelt) and the bird arrival date. Migratory waterfowl might not track the exact day that a specific resource becomes available but may display a lag effect, where birds arrive before (positive lag value) or after (negative lag value) the resource becomes available [15]. In order to obtain the focal lag (i.e., the number of days before or after the resource available or snow melt dates), we tested the predictive ability of the models across a series of temporal lags, by adding or subtracting days to the resource becoming available or the snow melt dates, following Laforge, Bonar [51]. For example, if a bird arrives at the stopover site two days after seed availability, the model with a lag of two (by subtracting two from the day to the seed availability date) would have the highest likelihood.

We executed RSFs in two regions for each species: the core stopover area Northeast China Plain and the breeding sites. We ran two models in the Northeast China Plain stopover sites (35°-55°N latitude) using vegetation and seed availability as the predictors, and three both at the mid-latitude (around 46°-49°N latitude) and arctic breeding sites (around 70°-75°N latitude) using vegetation availability, the onset of snowmelt, and the end of snowmelt as predictors. The response variable was the probability of bird arrival. The presence was set to 1 and absence was set to 0. For each of the five waterfowl species, within each region, we fit a mixed-effects conditional logistic regression model using a binomial distribution as the link function. The fixed effect for each model was a specific type of temporal lag, i.e., lag days to 1) the spring vegetation emergence, 2) the start of seed availability, 3) the onset of snowmelt, or 4) the end of snowmelt. For each individual tracked in each year, we obtained the identity of the track (track ID), named as individual (bird ID) and year (ID-year). The track ID was included in the model as a random effect. We used the R package glmmTMB [52] to run the analyses.

We modelled the lags in days to food availability or the snow melt conditions that fell between the 10th and 90th percentile of all locations within each region used by herbivorous waterfowl. We selected the top model for the focal lag based on the lowest Akaike Information Criterion (AIC) [51]. The range around the optimal lag was also estimated by predicting the day at which interpolated log-likelihood values have a difference of 1 (ΔAIC = 2). The difference in range was then calculated by the number of days between the minimum and maximum estimate. The predictive performance of the top models with the optimal estimated lag was evaluated by k-fold cross-validation [51, 53], in according to previous studies [54, 55]. This k-fold cross-validation procedure withheld a fraction of the data using a k-fold partitioning [54], where k represented the number of partitions (i.e., test-training sets), ranging from two to *N*-1 (number of observations minus one) [54, 55]. We split our origin samples by the identity of the track (track ID) into k (k = 3) training-testing sets and 10 bins. We trained our model iteratively on two out of the three datasets using a logistic regression. The remaining dataset was used for validation. To examine model performance, the pattern of predicted RSF scores for partitioned test-training sets is investigated against categories of RSF scores (bins) [55]. We calculated Spearman rank correlations (r_s_) between 10 RSF bin ranks and three test-training sets (k = 3). A model with good predictive performance would be expected to be one with a strong positive correlation (i.e., a high r_s_ value) [55]. For each region and each species, we therefore considered the predictor used in the model with the highest r_s_ value as the cue used by herbivorous waterfowl in timing their migration [56].

The calculation of the green wave was performed in MATLAB R2020a [57]. Calculating the timing of seed wave, investigating resource utilization, and analysing the relationship between resource availability and bird arrival were performed in R 4.1.0 [58].

## Results

### Resource utilization along the migration routes

We analyze the relationship between spatiotemporal bird migration patterns (Table S1 and Figure S3) and the availability of two food resources (seeds and newly emerged vegetation). Spring migration tracks (*n* = 167) from 2015 to 2021 of five waterfowl species within the East Asian – Australasian Flyway were used: 22 tracks of swan geese, 8 of tundra swans, 54 of tundra bean geese, 63 of greater white-fronted geese, and 20 of lesser white-fronted geese. Swan geese migrate around 2500 km from their wintering grounds in Poyang, China to reach their mid-latitude breeding sites in Inner Mongolia and Mongolia (Table S1 and Figure S3*C*). The other four species are arctic breeders, with a migration distance over 5000 km (Table S1 and Figure S3*A, B, D, E*). Seed resources at farmlands are all distributed below 52°N latitude (Figure S4). Tracked birds utilize both natural habitats (water and surrounding wetlands, grassland, and shrublands) and farmlands (especially in the core stopover sites in the Northeast China Plain). In Russia (above 52°N latitude), waterfowl mainly use water, shrublands, grasslands, and wetlands. They briefly stay at wetlands or rivers that occur within forests, which explains the high proportion of forest habitats used in this region (Figure S4*F-H*). At breeding sites, tracked birds use grasslands, wetlands, and shrublands as their main habitats (Figure S4).

### Herbivorous waterfowl track multiple resource waves during spring migration

Herbivorous waterfowl in East Asia adopt a strategy in which they use both the seed wave and the green wave during spring migration (Figure 2). All study species arrive after seeds become available (using the end of snowmelt as a surrogate) and depart after the new vegetation become available in their core stopover sites in the Northeast China Plain (Figure 2 and Figure S5). The period when seeds become available in this region is at the end of March (86 ± 31 Julian days, Figure 2), whereas the emergence of new vegetation occurs around early May (114 ± 30 Julian days, Figure 2). Birds arrive at the core stopover sites in early April (94 ± 14 Julian days, Figure 2). All arctic-breeding species depart from the Northeast China Plain region around mid-May (131 ± 10 Julian days, Figure 2E, *H, K, N*). All study species arrive at their breeding sites around the onset of vegetation growth (Figure 2). In the Russian stopover sites where they briefly pause, the observed arrival timings primarily match the end of snowmelt (around the end of May and the beginning of June). Birds reach their arctic breeding sites around early June (Table S1, Figure 2F, *I, L, O*, and Figure S5).

**Figure 2.**
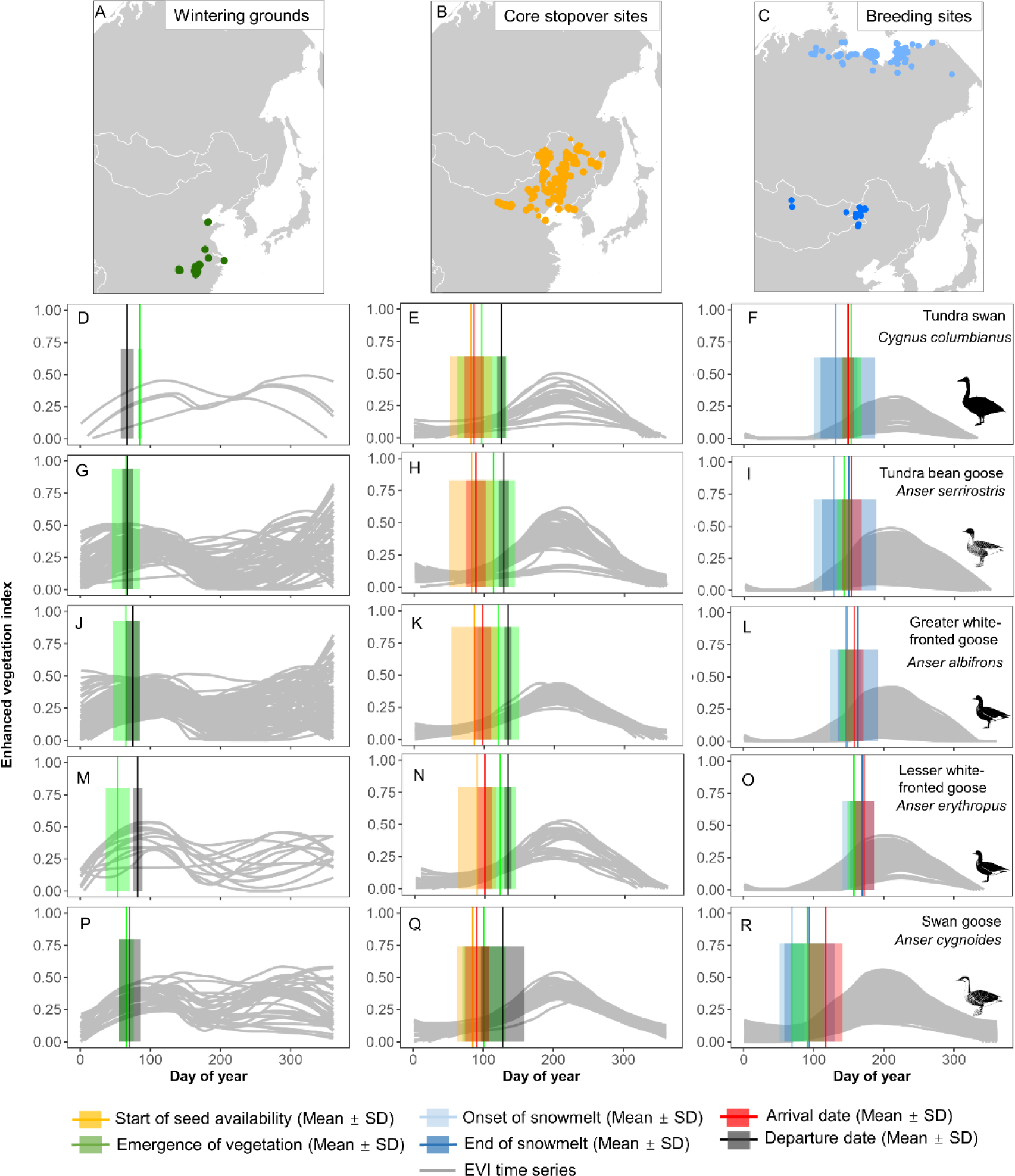
Tracking multiple resource waves during spring migration. All study species **(D-R)** arrive (vertical red line with error bar) at the Northeast China Plain after the start of seed availability (vertical orange line with error bar) and departed (vertical black line with error bar) from this region after the emergence of vegetation (vertical green line with error bar), thereby use both food resources. Greater white-fronted geese and lesser white-fronted geese depart from the wintering grounds **(J, M)** and arrive at the Northeast China Plain **(K, N)** later than larger-bodied species **(D, E, G, H, P, Q)** to better use the newly emerged vegetation (vertical green line with error bar). All study species arrive at the breeding sites around the time vegetation emerges (**F, I, L, O, R**). Grey lines show annual food availability indicated by enhanced vegetation index (EVI) time series. Dots show the centroid locations of wintering grounds (green dots in **A**), core stopover sites in the Northeast China Plain (orange dots in **B**), breeding sites in the mid-latitude (dark blue dots in **C**) and arctic (light blue dots in **C**) used by five herbivorous waterfowl.

Herbivorous waterfowl indeed spend relatively more time on farmland when they arrive at their core stopover sites, and then shift to spend more time on natural habitat after newly emerged vegetation becomes available (Figure 3, Figures S6 and S7). For all study species, the proportion of habitat use on farmlands at daytime is 42% before the emergence of vegetation, whereas this decreases to 32% after the emergence of vegetation, resulting in a 10% shift in foraging distribution from farmlands towards natural habitats (i.e., grasslands, shrublands, wetlands, and water areas) (Figure 3). This habitat shift pattern is more obviously in the two larger-bodied species, with a 28% shift for tundra swans and 16% for swan geese (Figure 3).

**Figure 3.**
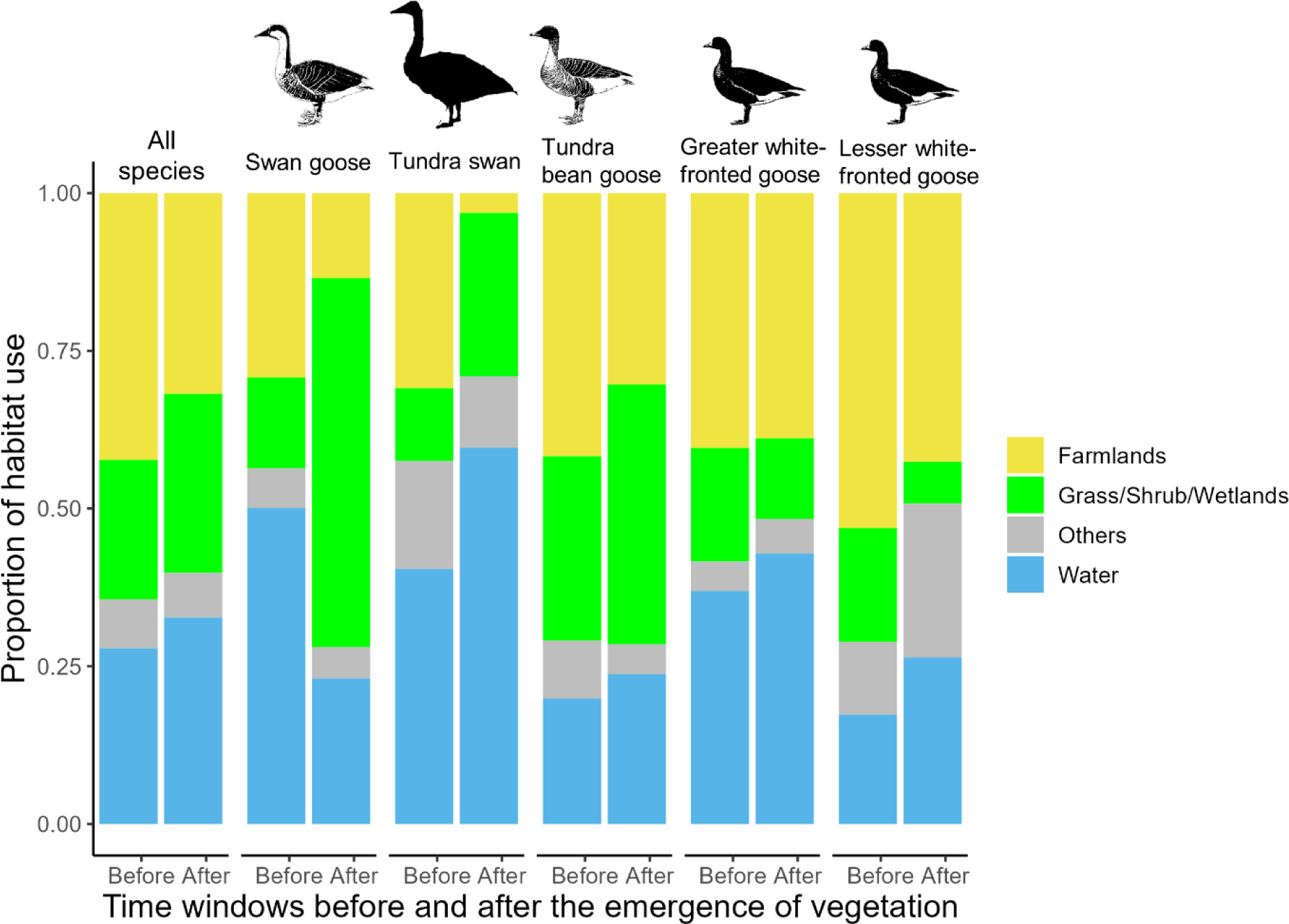
Foraging distribution before and after the emergence of vegetation. The proportion of habitat use showing foraging distribution at core stopover sites in the Northeast China Plain was calculated using the proportion of GPS locations during the daytime on each land cover type. The proportion of GPS locations during the daytime on farmlands decrease, shifting on average 10% towards natural habitats after the emergence of vegetation for all study species. Larger-bodied herbivorous waterfowl (i.e., tundra swan *Cygnus columbianus* and swan geese *Anser cygnoides*) have relatively more shifts from farmlands to natural habitats after the emergence of vegetation than relatively smaller-bodied species (i.e., tundra bean geese *A. serrirostris*, greater white-fronted geese *A. albifrons*, and lesser white-fronted geese *A. erythropus*).

### Variation in tracking multiple resource waves among species

Although all study species track multiple resource waves during spring migration, the specific spring migration timing varies slightly among species (Figure 2 and Figure S8). Species with a relatively larger body size (tundra swans and tundra bean geese) show a relatively earlier departure from the wintering sites (*F = 9.98*, *df* = 4, *P < 0.001*, Figure 2D, *G*, Figure S8*A,* and Table S1), and a relatively earlier arrival on the Northeast China Plain (*F = 20.08*, *df* = 4, *P < 0.001*, Figure 2E, *H*, Figure S8*C* and Table S1), in comparison to smaller-bodied species (greater white-fronted geese and lesser white-fronted geese; Figure 2J, *K, M, N*). The two smaller-bodied geese stay in the wintering grounds for an extended period of time after the emergence of vegetation: 10 days in greater white-fronted geese and 28 days in lesser white-fronted geese (Figure 2J, *M*). The departure time from the core stopover sites is significantly earlier for mid-latitude breeding swan geese than the other four arctic-breeding species (*F = 10.45*, *df* = 4, *P < 0.001*, Figure 2, Figure S8*D* and Table S1). The smallest-bodied lesser white-fronted geese arrive at the arctic breeding sites the latest among all five herbivorous waterfowl species (*F = 40.67*, *df* = 4, *P < 0.001*, Figure 2, Figure S8*B* and Table S1).

The arrival probability at the core stopover sites and breeding sites in relation to the availability of food waves and snow conditions shows that optimal lags to the availability of multiple resources also vary slightly among species (Figure 4 and Table 1). For arctic-breeding species, the highest arrival probability occurs after the start of seed availability and before the emergence of vegetation in the core stopover sites (Figure 4A, *C, E, G*). Tundra swans arrive at the core stopover sites soon (only one day) after seeds become accessible (Table 1 and Figure 4A), whereas smaller-bodied goose species arrive later, after seeds have already been available for about one week (for tundra bean geese; Figure 4C) to up to three weeks (for greater white-fronted geese and lesser white-fronted geese; Figure 4E and G). On the arctic breeding grounds, three goose species (tundra bean geese, greater white-fronted geese, and lesser white-fronted geese) arrive at their breeding sites after the plant emergence (Table 1, Figure 4D, *F*, and *H*), whereas tundra swans arrive at the breeding sites relatively earlier, slightly before the emergence of vegetation (Table 1, Figure 4B). Mid-latitude breeding swan geese arrive at the core stopover sites and the breeding sites around three weeks after the local onset of vegetation growth (Table 1, Figure 4I and J).

**Figure 4.**
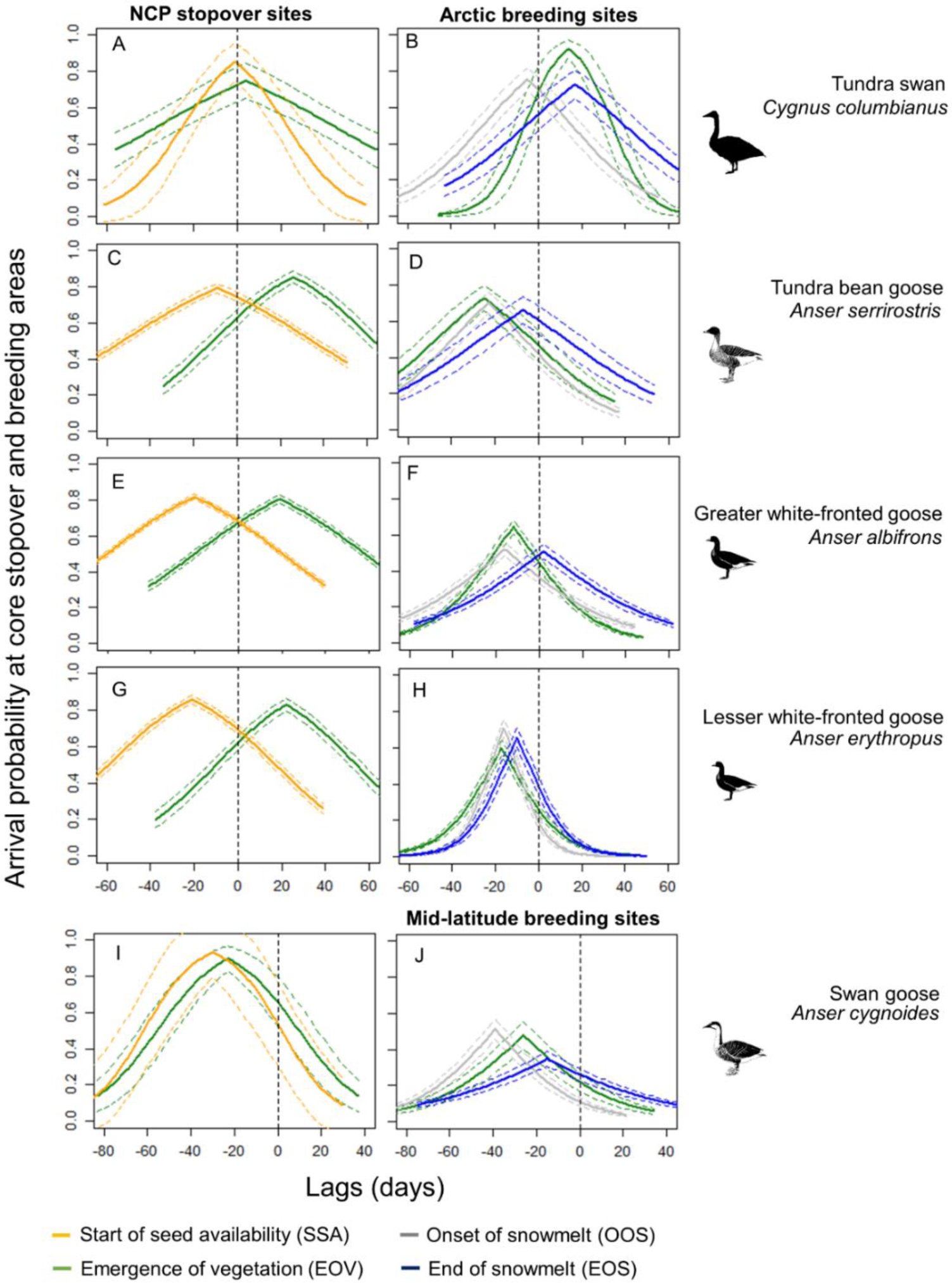
Top resource selection models showing the species variation in the use of multiple resources. Arrival probability in relation to the availability of seeds and newly emerged vegetation for East Asian migratory herbivorous waterfowl during spring migration. The highest arrival probability for arctic-breeding birds occurred after the start of seed availability (SSA; orange lines) and before the emergence of vegetation (EOV; green lines) in the Northeast China Plain (NCP) stopover sites **(A, C, E, G)**. All arctic-breeding geese reached their breeding sites after the emergence of vegetation (EOV in **D, F, H**; green lines) and close to the end of snowmelt (EOS in **D, F, H**; blue lines). Tundra swans arrived at the arctic breeding ground earlier (after the onset of snowmelt; OOS in B; grey lines). Mid-latitude breeding swan geese **(I, J)** reached the Northeast China Plain and the breeding sites after the emergence of vegetation (EOV; green lines). Colored dashed lines represent 95% confidence intervals. Vertical dashed lines represent the bird arrival date. Lags (x axis) indicate the lag days in comparison to the timing of resource availability or the timing of snowmelt.

**Table 1.** Top resource selection models showing that species using multiple resource waves in migration but relying on specific cues in specific region. The estimated optimal lags (days) on resource availability and snowmelt conditions shows the variation in tracking multiple resource waves at core stopover sites the Northeast China Plain and the breeding sites for five East Asian herbivorous waterfowl species. The estimated range represents the day at which interpolated log-likelihood values have a difference of 1 (ΔAIC = 2). The difference is the number of days between the minimum and maximum estimate of the range. Bold font represents strongest models evaluated by the relatively stronger k-fold cross-validation score (r_s_) in each region and for each species.

| Region | Species | Models | Estimated days with range | Difference | Log-likelihood | K-fold scores ($r_s$) | |
| --- | --- | --- | --- | --- | --- | --- | --- |
|  |  |  |  |  |  | Mean | SE |
| Northeast China Plain | Tundra swan | Plant * | 4 (-3.4, 13.7) | 17.1 | -213.57 | 0.427 | 0.450 |
|  |  | Seed † | **-1 (-5.5, 4.8)** | **10.3** | **-24.10** | **0.571** | **0.169** |
|  | Tundra bean goose | Plant | **26 (24.3, 27.8)** | **3.5** | **-101.14** | **0.873** | **0.125** |
|  |  | Seed | -9 (-11.1, -5.9) | 5.2 | -42.04 | 0.503 | 0.143 |
|  | Greater white-fronted goose | Plant | **19 (17.7, 21.7)** | **4** | **-1522.14** | **0.935** | **0.025** |
|  |  | Seed | -20 (-20.7, -18.9) | 1.8 | -4067.08 | 0.394 | 0.278 |
|  | Lesser white-fronted goose | Plant | **22 (20.2, 24.2)** | **4** | **-466.73** | **0.914** | **0.014** |
|  |  | Seed | -21 (-22.2, -19.9) | 2.3 | -1398.77 | 0.358 | 0.053 |
|  | Swan goose | Plant | -23 (-25.1, -19.5) | 5.6 | -1107.01 | 0.403 | 0.397 |
|  |  | Seed | **-30 (-35.7, -20.6)** | **15.1** | **-1380.99** | **0.487** | **0.546** |
| Breeding sites | Tundra swan | Plant | **14 (12.9, 16)** | **3.1** | **-6198.94** | **0.530** | **0.130** |
|  |  | Onset of snowmelt | -5 (-6.6, -2.3) | 4.3 | -5916.80 | 0.417 | 0.451 |
|  |  | End of snowmelt | 17 (16.1, 17.4) | 1.3 | -6789.48 | 0.523 | 0.108 |
|  | Tundra bean goose | Plant | -25 (-26, -23.5) | 2.5 | -336.80 | 0.301 | 0.164 |
|  |  | Onset of snowmelt | -23 (-23.4, -21.6) | 1.8 | -484.11 | 0.253 | 0.165 |
|  |  | End of snowmelt | **-7 (-9.5, -4.7)** | **4.8** | **-514.02** | **0.572** | **0.152** |
|  | Greater white-fronted goose | Plant | -12 (-12.2, -11) | 1.2 | -6332.97 | 0.628 | 0.182 |
|  |  | Onset of snowmelt | -16 (-17.1, -15.6) | 1.5 | -7455.17 | 0.434 | 0.081 |
|  |  | End of snowmelt | **2 (0.5, 3.1)** | **2.6** | **-7569.63** | **0.859** | **0.093** |
|  | Lesser white-fronted goose | Plant | -17 (-17.8, -16.9) | 0.9 | -2464.9 | 0.406 | 0.140 |
|  |  | Onset of snowmelt | -16 (-16.9, -15.9) | 1 | -1999.08 | 0.384 | 0.082 |
|  |  | End of snowmelt | **-10 (-10.1, -9.9)** | **0.2** | **-2171.43** | **0.648** | **0.128** |
|  | Swan goose | Plant | **-26 (-26.3, -25.8)** | **0.5** | **-3562.83** | **0.595** | **0.340** |
|  |  | Onset of snowmelt | -39 (-39.7, -37.7) | 2 | -3449.32 | 0.312 | 0.730 |
|  |  | End of snowmelt | -15 (-16, -14.4) | 1.6 | -3785.92 | 0.466 | 0.242 |

\* The model using the spring vegetation emergence as a predictor
† The model using the start of seed availability as a predictor

All study species use multiple resource waves during spring migration, but variation among species is observed, as indicated by the best resource selection models evaluated by k-fold cross validation (higher score, stronger effect; Table 1). At the core stopover sites, the models for larger-bodied species (tundra swans and swan geese) using the start of seed availability as a predictor have relatively higher k-fold cross-validation scores, indicating the use of the seed wave as a cue. For smaller-bodied arctic-breeding geese, the models using the emergence of plants as a predictor have a stronger effect, indicating the use of the green wave as a cue. At the breeding sites, the models for larger-bodied species using the emergence of vegetation have the highest k-fold cross-validation scores, suggesting these waterfowl species track the green wave. For the relatively smaller-bodied species (tundra bean geese, greater white-fronted geese, and lesser white-fronted geese), the models that use the end of snowmelt as predictors show the strongest effect, indicating the use of the snow-free time as a cue.

## Discussion

We test a new migration strategy in which East Asian herbivorous waterfowl use multiple resource waves to time their spring migration. The variation in tracking multiple resource waves for each species and in each region are identified regarding the use of the seed wave (agricultural seeds in farmland) and the green wave (newly emerging vegetation in natural habitat). Larger-bodied species follow the seed wave and smaller-bodied ones adopt the green wave as the cue at the core stopover sites. When arriving at the breeding sites, larger-bodied species track the green wave whereas smaller-bodied ones track the snow-free time. These strategies in which migration is timed to be able to take advantage of both natural and anthropogenic food resources allows East Asian herbivorous waterfowl to exploit the vast and variable landscape along their migration routes.

Consistent with our first prediction, East Asian herbivorous waterfowl time their staging in the core stopover sites Northeast China Plain to make use of both anthropogenic and natural food resources. Seeds left over from previous year’s harvest on the unfrozen ground (from the beginning of April) and newly emerged plants (in early May) allow migratory birds to replenish their energy for an extended period of time during stopover, which facilitates them arriving at the breeding grounds in an optimal condition. The Northeast China Plain contains 15% of the total farming area in China, with maize, soybean, and rice being the three major crop types [59]. Crop seeds on farmlands provide a high-nutrient food source for herbivorous waterfowl [28], especially during early spring, when snow just starts melting and food in the natural habitat is insufficient. Mechanical ploughing (Figure S2*A*) at the end of April predicts sowing (Figure S2*B* and *C*). In early May, seeds covered with pesticides are sowed (Figure S2*C*), which means crop seeds on farmlands are no longer useable for waterfowl (personal field observations and personal communications with local farmers; Figure S2). When seeds are not available anymore, all study species show a pattern in which they shift towards the use of the natural habitat.

The exact migration timing differs among species relating to their body size. One explanation could be that larger-bodied species do not rely on the emergence of natural food resources as much as the smaller-bodied goose species. Larger-bodied arctic-breeders generally invest more time on breeding activities and face a more time-constrained window in which they have to lay eggs, molt, and raise chicks, than smaller-bodied species [4, 60, 61]. The slower pace in their life-histories (i.e., laying larger eggs with a longer incubation and gosling-rearing period) drives birds to reach the breeding grounds and lay eggs earlier [61]. This is particularly the case for tundra swans, with the largest body size among the five herbivorous waterfowl studied, which arrive at the core stopover sites and the arctic breeding sites the earliest and overtake the green wave, but which can use underwater and underground food resources once the ice melts [62]. Smaller-bodied grazing species, namely the greater and lesser white-fronted geese, depart later from the wintering site (Figure 2J, *M* and Table S1), so that they can forage there on newly emerging grasses before moving north [63–65]. They thereby arrive much later in the core stopover sites and the breeding sites. This strategy enables them to maximize their time foraging on the newly emergent shoots in their wintering and stopover areas [8]. Tundra bean geese, slightly larger than greater and lesser white-fronted geese, arrive at the core stopover sites only one week after seeds have become available, spend longer stopovers to make use of both resources (Table S1), and arrive at the breeding sites after the emergence of vegetation. These patterns showing species-dependent resource utilization at different times likely explain how the species can coexist in a shared landscape.

In line with our second prediction, larger-bodied herbivorous waterfowl, namely tundra swans and swan geese, use the seed wave to time their arrival at the core stopover sites. They generally are less sensitive to forage quality, and therefore can utilize a relatively more diverse assortment of food resources [39, 64]. They prefer open water where underwater food resources can be easily accessed (e.g., tubers and algae) that, just like seeds, only become accessible after snowmelt [62]. This is in line with the habitat use after they reach the stopover sites, that they spend relatively longer time in water than other smaller-bodied species (Figure 3). Tundra swan then follow the green wave to arrive at their arctic breeding sites the earliest. This provides direct benefits for larger-bodied waterfowl, such as prioritizing resource consumption, and having sufficient time to raise their offspring [61].

Different from our prediction, we found that three smaller-bodied arctic-breeding geese rely relatively more on natural food resources, and accordingly follow the green wave at core stopover sites. They arrive in the arctic after vegetation has newly emerged, using the snow-free time as a cue. While newly emergent vegetation provides a nutritional resource, melting snow facilitates the emergence of vegetation and provide a cue for animals to exploit resources at the correct time, which helps to reduce movement and foraging costs related to snow cover [51]. This finding is in line with a previous study reporting that East Asian geese arrive at their arctic breeding sites prior to the end of snowmelt [66]. We highlight that smaller-bodied goose species use the newly emerged vegetation as a primary cue to enable accumulating sufficient fuel during migration, and avoiding that they arrive too early and deplete their body stores while awaiting the local thaw, and at the meantime ensuring that their goslings benefit from abundant food resources later in the season [4, 6, 66].

Our study provides the first evidence for multiple resource tracking in waterfowl spring migration. The central tenet of resource tracking theory is that mobile consumers can benefit by moving to exploit phenological variation in resources across space [5]. Tracking multiple resource waves allows migratory herbivorous waterfowl to prolong the overall resource availability of natural and anthropogenic resources. Similar to our results, previous study found that forest birds and stream fishes increase foraging opportunities by feeding on emerging stream insects early in the year and then switching to terrestrial arthropods later on, in response to the asynchronous occurrence of fluxes of invertebrates between terrestrial and freshwater environments [67]. Recent empirical work shows that elephants track both vegetation green-up in natural habitat and farmland brown down in a coupled human-nature system [68], maximizing their energy intake by integrating agricultural food resources when natural food resources is limited. To further our understanding on the response of species to environmental change, the capability of tracking multiple resource waves should be further tested in additional study systems and different geographical regions.

### Conservation implications

Climate change might induce shifts in vegetation phenology in the spring [69–71], thereby changing food availability for herbivorous waterfowl at a different pace at different sites [72]. A reduced predictability of food supply due to phenological asynchrony would be exacerbated when different food resources are exploited. As a consequence, species may struggle to advance their migration timing to match food availability [73]. Even though some goose species can accelerate their migration speed under global warming, they might still suffer population declines if they cannot store sufficient nutrients for egg-laying [60]. For species taking advantages of the seed wave, migration timing and pace could be influenced by changes in cultivation practices [36]. For example, changes in farming calendars might affect seed availability [17, 23, 36]. Different responses to environmental change between the timing of cultivation and bird migration phenology might cause ecological traps and population declines of migratory birds [74, 75]. Therefore, changes in habitat management should consider conservation needs, especially for species that have come to depend on anthropogenic food resources [76].

### Study limitations

We use two different seasonal metrics derived from satellite imagery (i.e., timing of snowmelt and the second derivative of EVI) to measure two different resource waves (i.e., the start of seed availability and the emergence of vegetation). Given that data that directly measure food availability site-by-site in the field are costly to obtain, satellite-derived indices provide an effective way to measure the seed wave and green wave. They cover more detailed pixel-by-pixel information in space and time, and better match the GPS locations for tracked birds. However, satellite-derived indices necessarily contain uncertainties. Firstly, our estimated plant emergence time (the green wave) based on EVI time-series data reflects the time that the greening increases at the fastest rate. This time might be later than the shoot emergence time. Secondly, since seeds are generally spilled and available during the previous fall and carried over into spring after soil thawing, it is crucial to investigate how many seeds are actually available in spring, and how much of those seeds are eaten by waterfowl moving through. Ground-truth data can further optimize the derived plant and seed availability.

## Conclusions

We test and demonstrate how migratory herbivorous waterfowl time their spring migration using both the post-harvest seed availability in cultivated landscapes and the emergence of vegetation in more natural landscapes. Specifically, we document how the dynamics of anthropogenic and natural resources act in combination to shape the spring migration phenology of herbivorous waterfowl. To what extent populations of migratory herbivorous waterfowl benefit from or are threatened by this migration strategy of tracking multiple resource waves needs to be thoroughly investigated. For example, how the seed wave itself is affected by climate and agricultural practices, is still unknown. A proper understanding of the impact of global environmental change and different human cultivating activities on migratory bird species will guide effective habitat management for migratory bird conservation worldwide.

## Acknowledgments

We thank Kun Xiao (Sun Yat-sen University, China) for helping processing the remote sensing data. We are grateful for John Takekawa, Sivananintha Balachandran, Guanhua Liu, Hao Luo, Duanji Tao, Saras Kumar Behera, Murugesan Ponnaian, Wenyuan Zhang, Yanjie Xu, Boyu Gao, Zhiyuan Lv, Lingtong Yang, Yongzhi Xiao, and Yunrui Liu who joined the field work. This research was supported by the National Natural Science Foundation of China (Grant No. 42301055, No. 41471347, and No. U22A20563).

## Author contributions

Conceptualization, F.X., P.G., and Y.S.; Data curation, F.X., J.W., G.L., J.L., B.W., W.Z.W., and Y.S.; Methodology, F.X., W.W., J.W., Q.X., W.Z.W., and Y.S.; Software, F.X., W.W., and J.W.; Investigation, F.X., J.W., and Y.S.; Formal Analysis, F.X., and W.W.; Visualization, F.X., and Y.S., Writing—Original Draft: F.X., and Y.S.; Writing—Review & Editing: F.X., B.W., F.A.L., Z.M., Y.Y., P.G., B.X., and Y.S.; Supervision, B.W., F.A.L., Z.M., Y.Y., and Y.S.; Project Administration, F.X., and Y.S.; Funding Acquisition, F.X., G.L., and Y.S.

## Declaration of interests

The authors declare that they have no competing interests.

## Data and materials availability

Movement tracks of five herbivorous waterfowl species are stored in Movebank (www.movebank.org), under ID 731644623 for lesser white-fronted geese, 1614083125 for greater white-fronted geese, 1614098614 for tundra bean geese, 1614096661 for swan geese, and 1614102131 for tundra swans. The authors declare that all other data supporting the findings of this study are available in the main text or the supplementary materials.

## Supporting information

Supplementary information is available for this paper.

## Notes

### Competing Interest Statement

The authors have declared no competing interest.

### Summary of Updates

Abstract and Figure 2 revised.

