## Supplementary Information for "Migratory herbivorous waterfowl track multiple resource waves during spring migration"

**This PDF file includes:**

Supplementary Text
Figures. S1 to S10
Tables S1
References (1 to 9)

**Supplementary Text**

Identifying stopover sites using the space-time permutation model

The space-time permutation model [1] is defined by a cylindrical window with a circular base representing space and height representing time. The cylinder starts with a single point and increases its base and height until it reaches a maximum value. The maximum value for the base and height is selected based on the migration behavior of targeted species. In our case the maximum radius for the base was an empirical estimation of the longest distance birds might travel during their stay at the stopover sites, and for the height this was the potential length of stay at these sites. We calculated the expected number of GPS locations based on complete spatial randomness, and measured a Poisson generalized likelihood ratio using the expected and observed number of GPS locations to determine whether the cylinder contained a cluster or not [1]. The significant level of the identified clusters was evaluated by 999 Mont Carlo simulation, through comparing the rank of the maximum likelihood from the observed dataset with the random dataset.

For the best performance, the maximum spatial scanning window should not exceed 50% of all GPS locations, and the temporal window should be set to no more than 50% of the study period [2]. We thereby chose 50 km as the maximal spatial scanning window as the maximum foraging flight distance for geese are generally smaller than this [2-4]. Considering birds need to stay in stopover sites for at least 48 hour to settle and refuel [5], a minimum temporal scanning window of two days and a maximum temporal window covering 50% of the spring migration period were selected. GPS locations record in flight (with a speed > 1 km/hour) were excluded from the analysis.

Calculating spring vegetation emergence time

The Moderate Resolution Imaging Spectroradiometer (MODIS) Terra 8-day 250 m surface reflectance product MOD09Q1 was used to derive the spring vegetation emergence time. Data was downloaded from http://e4ftl01.cr.usgs.gov. The spatial range of the imagery covers the whole migratory flyway and the time period covers 2015-2021. A two-band Enhanced Vegetation Index (EVI2) was first calculated according to equation (1):

$$\boldsymbol{EVI2 = 2.5\frac{N - R}{N + \left(6 - \frac{7.5}{2.08}\right)R + 1}} \qquad (\mathbf{1})$$

Where N and R represent surface reflectance in near-infrared and red bands [6]. Cloud-free pixels were extracted in the EVI2 time series according to the band of QA (data product quality assessment). A method of moving median was used to interpolate the removed pixels with clouds. A daily EVI2 time series was created by interpolating missing data using spline interpolation. A Savitzky-Golay filter was applied for smoothing the time series and a logistic curve fitting function was used to fit the green-up phase in the reconstructed daily EVI2 time series [7, 8]. The spring vegetation emergence time was then determined by the date that reached the first local maximum value of the second derivative of logistic-fitted EVI curve.

Extracting pure pixels in the Northeast China Plain

It is challenging to extract the green wave from satellite imagery compared to field observations, because different smoothing methods and retrieval algorithms used to process satellite time series data for plant phenology could result in spatial variance in vegetation dynamics [7].

Moreover, plant phenology varies among different land cover types and mixed-pixel effect can negatively affect green wave estimation [9]. This is especially the case in the Northeast China Plain, where small and fragmented herbaceous areas are enveloped by agricultural fields, and in Eastern Siberian lowland, where natural wetlands and grasslands are embedded in forests. Nevertheless, we have taken this complication into account by only using pure pixels in the natural habitat in the Northeast China Plain, to lower the impact of the mixed-pixel effect.

In China, especially in the focal stopover sites in the Northeast China Plain, the natural habitat of migratory herbivorous waterfowl is fragmented (Figure S9*A*). As a consequence, the onset of plant growth derived from satellite imagery could be biased due to the mixed-pixel effect (e.g., small and fragmented herbaceous areas embedded in large agricultural fields). Moreover, it remains hard to extract the onset of new plant growth for the narrow belt grasslands surrounding wetlands if pixels are mixed with agricultural land. In order to remove the mixed-pixel effect and catch the green wave at natural habitats, we extracted pure pixels of spring vegetation emergence layer by overlaying both the 30 m land cover dataset of China and the 250 m grid of spring vegetation emergence layer derived from smoothed EVI time series. We calculated the centroids of the pixels of the 250 m spring vegetation emergence layer and created a 125 m buffer. Then we summarized the proportion of each 30 m land cover type within the buffer, Pi,

$$Pi = ni/N$$

Where ni is the number of pixels in the specific land cover type and N is the number of all pixels in the buffer. Then we selected pixels with values of Pi larger than 0.5 as pure pixels and thus obtained the corresponding land cover type in this pure pixel (Figure S9*B*).

**Figure S1.**

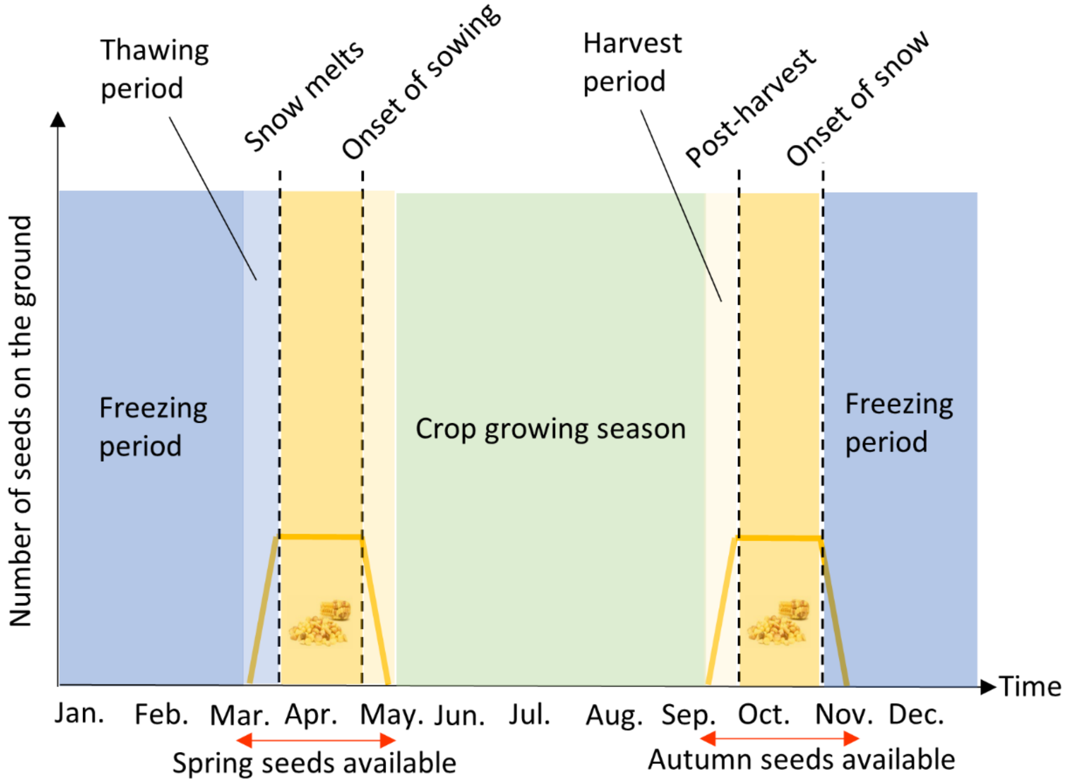


Schematic diagram of how the seed wave works. Seed availability period during the spring starts from the end of snowmelt (at the end of March) and ends at onset of sowing (in early May).

**Figure S2.**

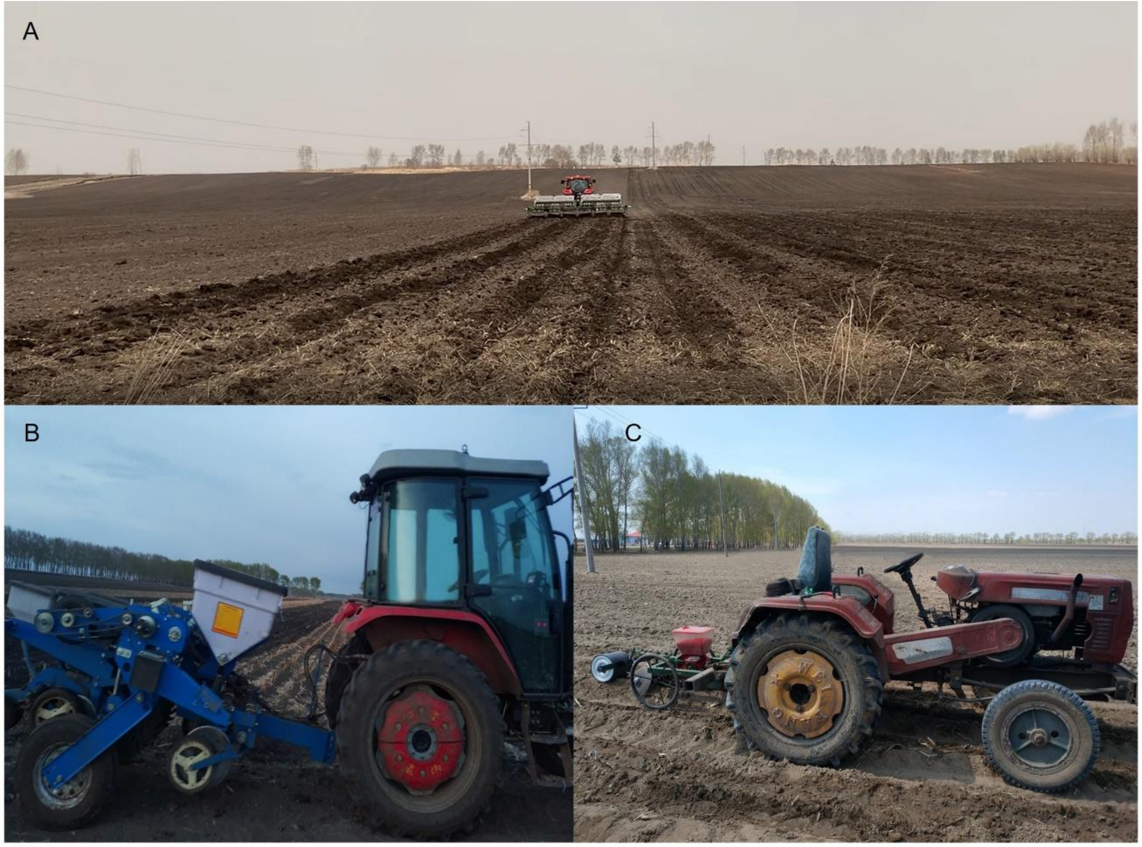


Human cultivation activities at a core stopover site in the Northeast China Plain. (A) Local farmers plough the farmland at Shuangshan Village, Nenjiang, Heihe on April 27th, 2021 (photo credit: Lingtong Yang). (B) Mechanical sowing of maize at Halahai town, Longjiang County, Qiqihar on May 1st, 2021 (photo credit: Anonymous). (C) Maize mixed with pesticides is sowed at farmlands located at Momoge, Zhenlai County, Baicheng on May 1st, 2021 (photo credit: Anonymous).

**Figure S3.**

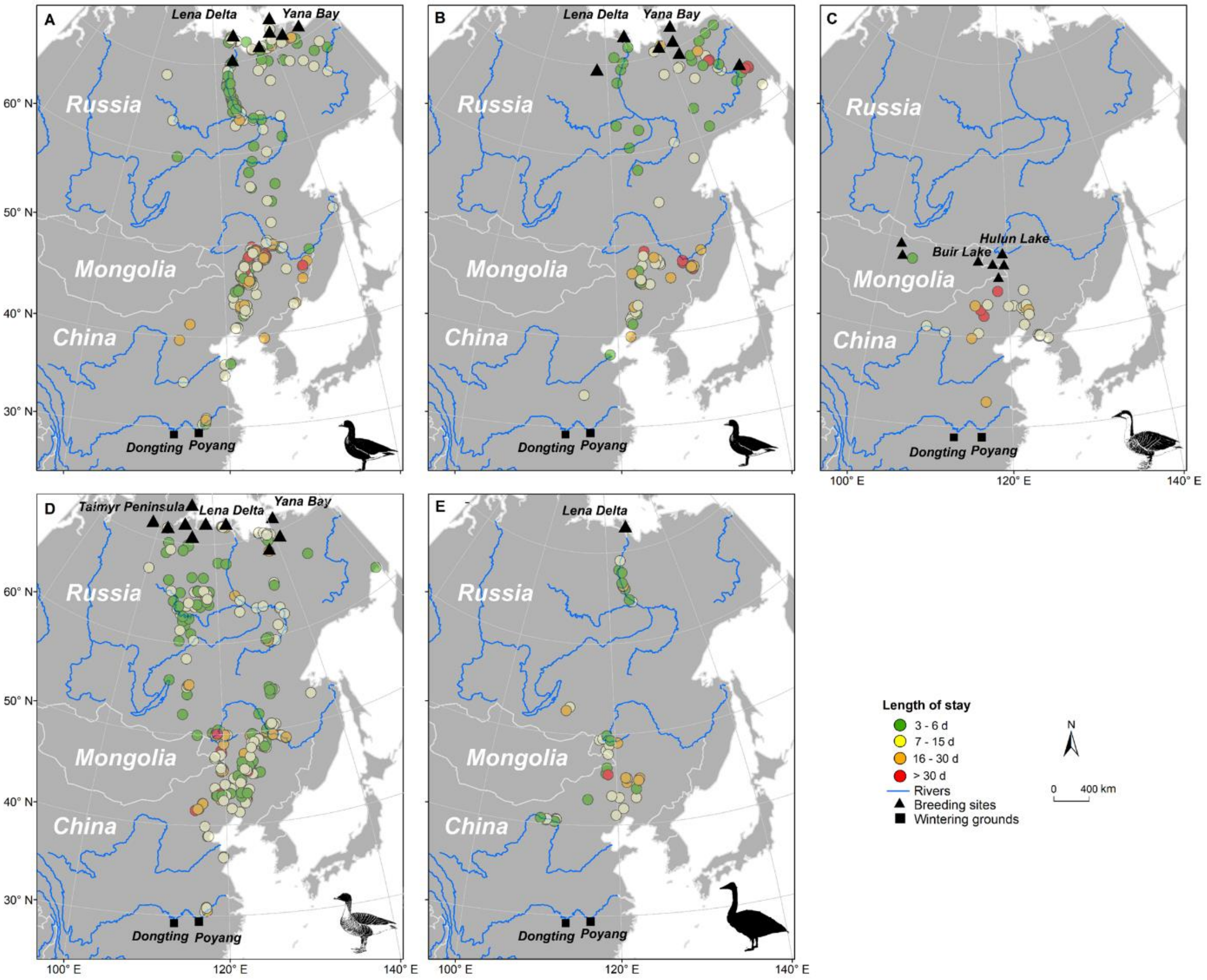


Stopover sites and the length of stay in each stopover site for A. greater white-fronted goose *Anser albifrons*, B. lesser white-fronted goose *A. erythropus*, C. swan goose *A. cygnoides*, D. tundra bean goose *A. serrirostris*, and E. tundra swan *Cygnus columbianus*.

**Figure S4.**

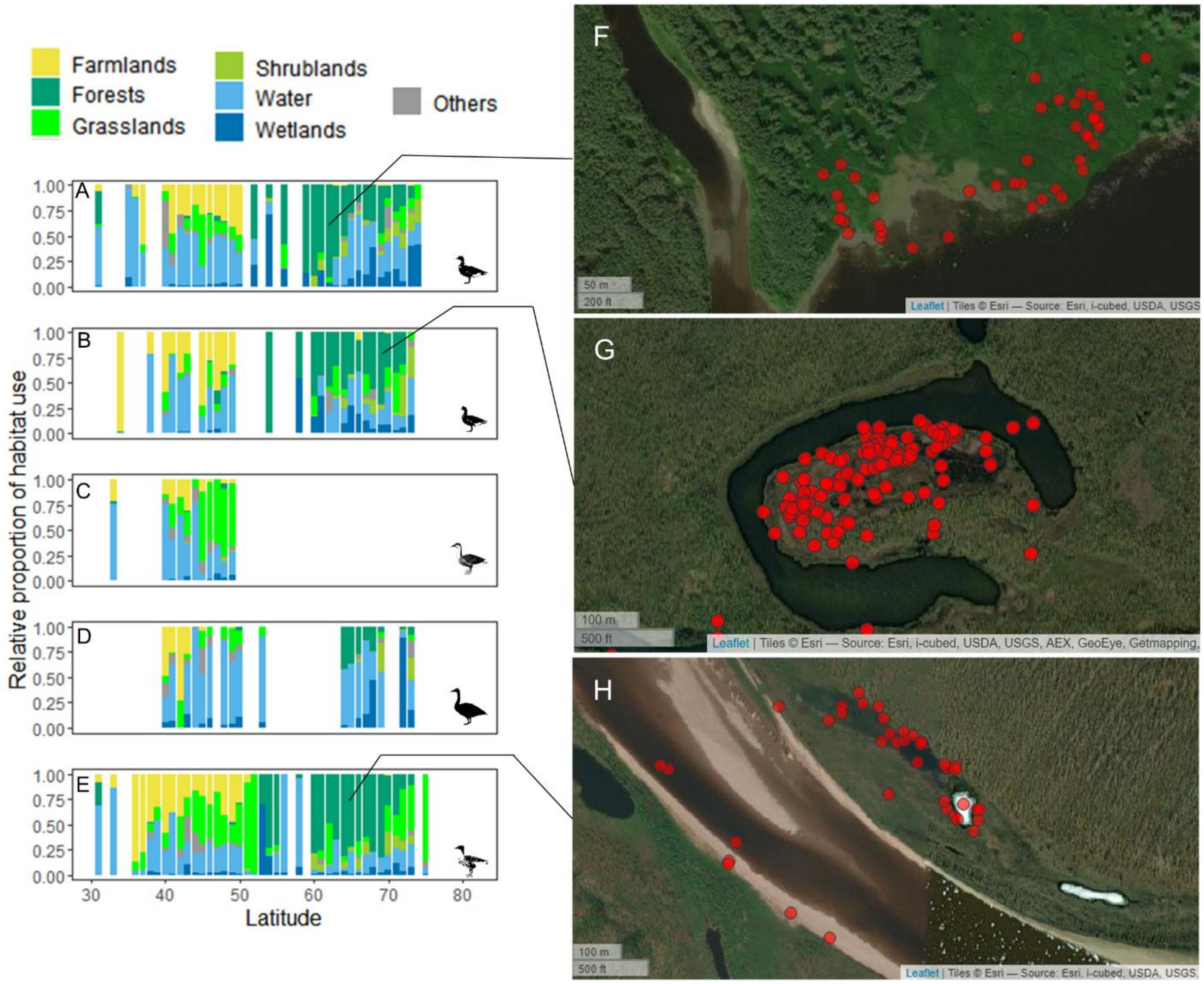


Proportion of habitat use along the spring migration route for A. greater white-fronted goose *Anser albifrons*, B. lesser white-fronted goose *A. erythropus*, C. swan goose *A. cygnoides*, D. tundra swan *Cygnus columbianus*, and E. tundra bean goose *A. serrirostris*. (F-H) GPS locations in the wetlands and water areas around forests explain relatively high proportion use of forests in Russian stopover sites.

**Figure S5.**

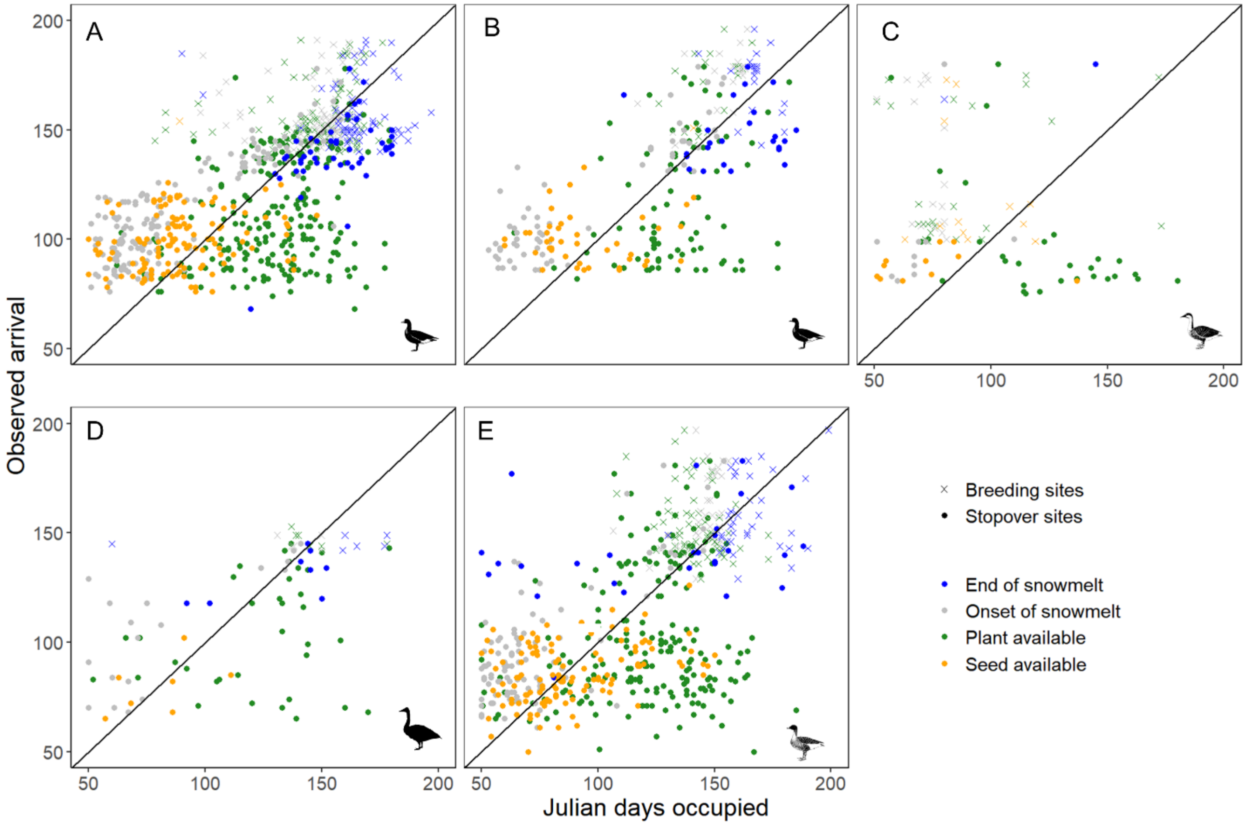


The relationship between observed arrival timing on the stopover sites and breeding sites with the mean value of different resource available date at each site for A. greater white-fronted goose *Anser albifrons*, B. lesser white-fronted goose *A. erythropus*, C. swan goose *A. cygnoides*, D. tundra swan *Cygnus columbianus*, and E. tundra bean goose *A. serrirostris*.

**Figure S6.**

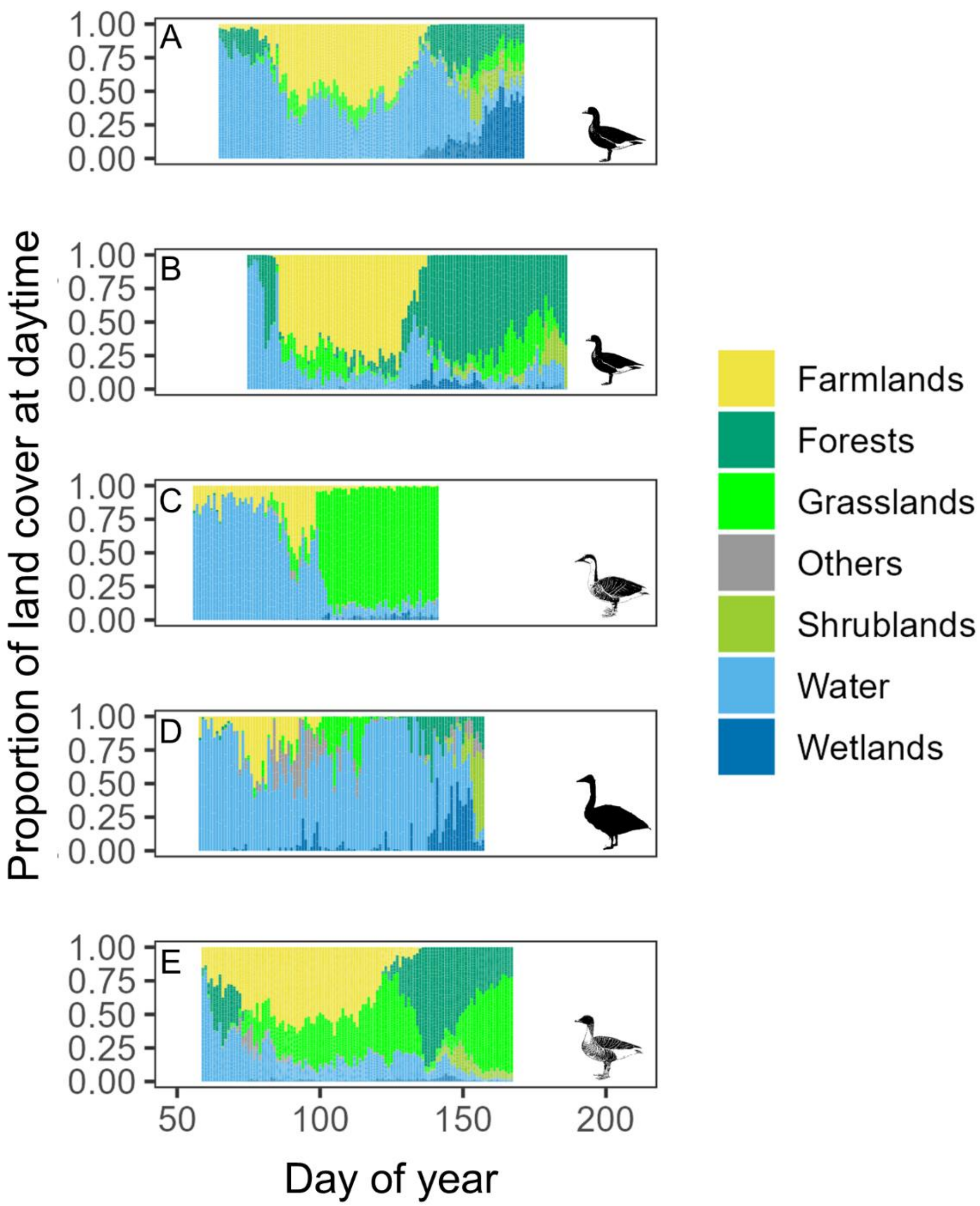


Habitat use at daytime during the whole spring migration period for A. greater white-fronted goose *Anser albifrons*, B. lesser white-fronted goose *A. erythropus*, C. swan goose *A. cygnoides*, D. tundra swan *Cygnus columbianus*, and E. tundra bean goose *A. serrirostris*.

**Figure S7.**

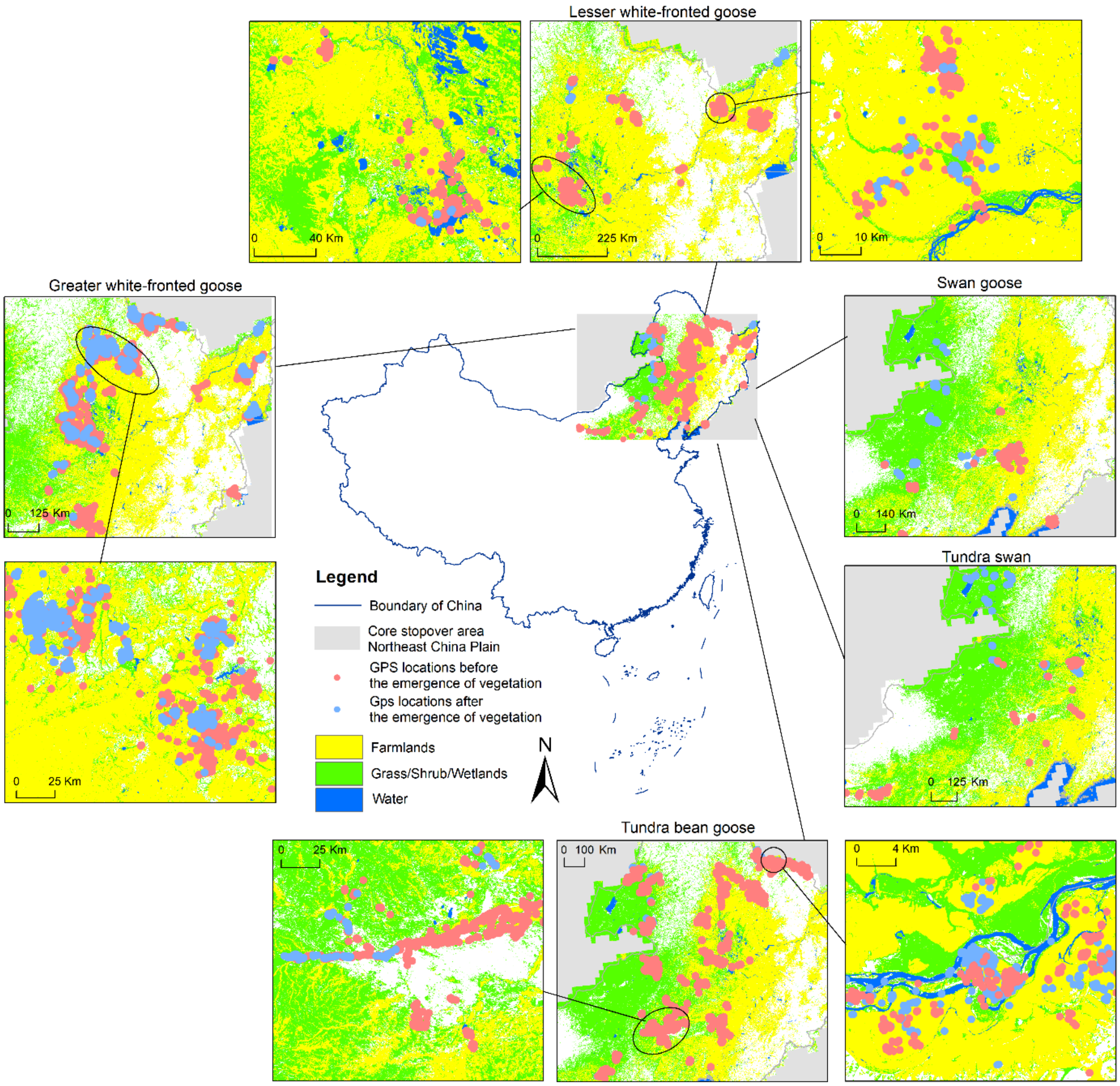


Foraging distribution on two types of food resources for herbivorous waterfowl in the core stopover area Northeast China Plain. Waterfowl use seeds on farmlands (yellow) and newly emerged plants on natural habitats (green). Dots showing bird GPS locations during the daytime before (pink) and after (shallow blue) the emergence of vegetation.

**Figure S8.**

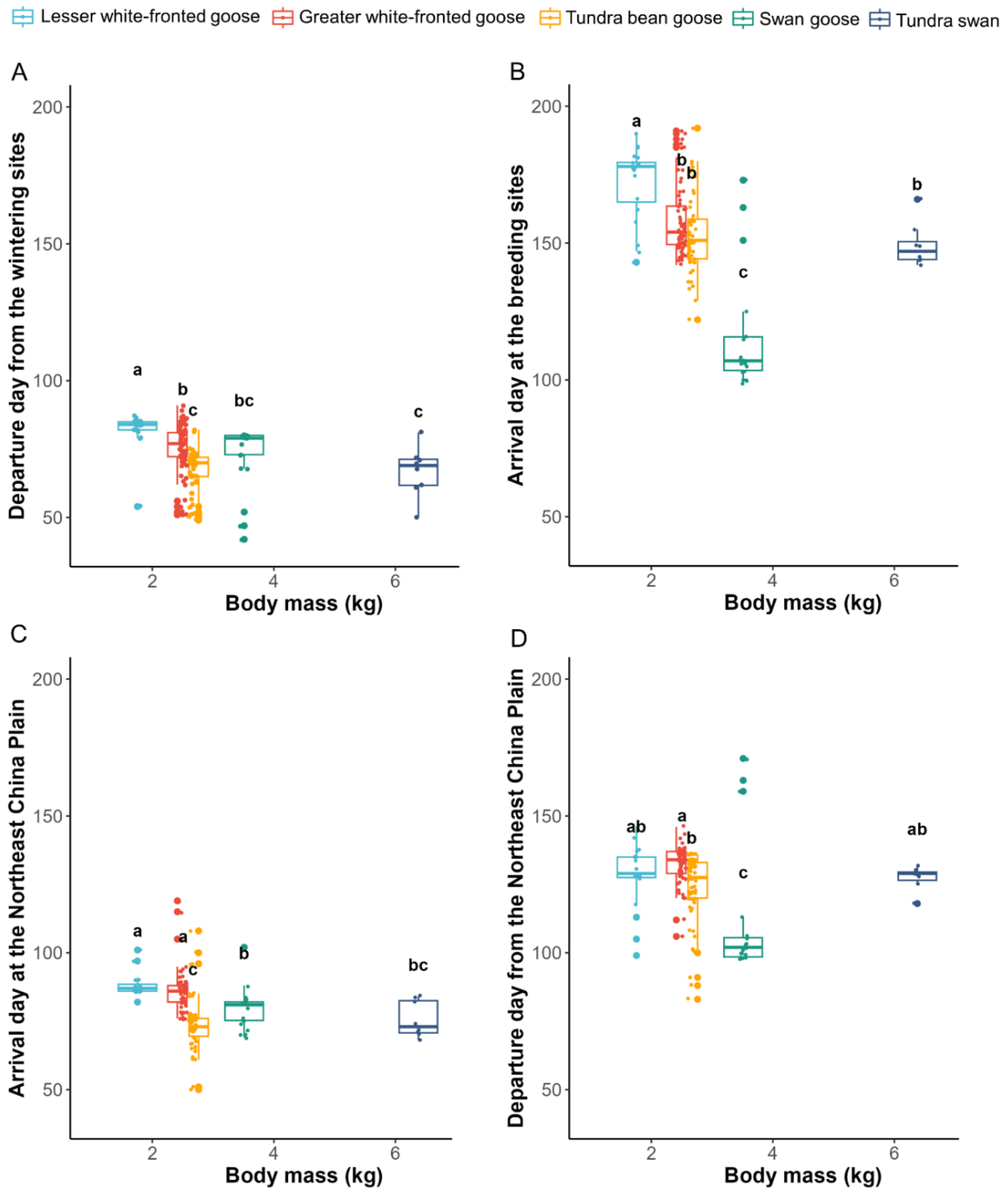


The relationship between migration timing and body size**.** Larger-bodied herbivorous waterfowl (A) depart earlier from the wintering grounds, (C) arrive earlier at core stopover sites Northeast China Plain, and (B) arrive earlier at breeding sites than smaller-bodied species. (D) Mid-latitude breeding swan geese depart earlier than arctic-breeding species.

**Figure S9.**

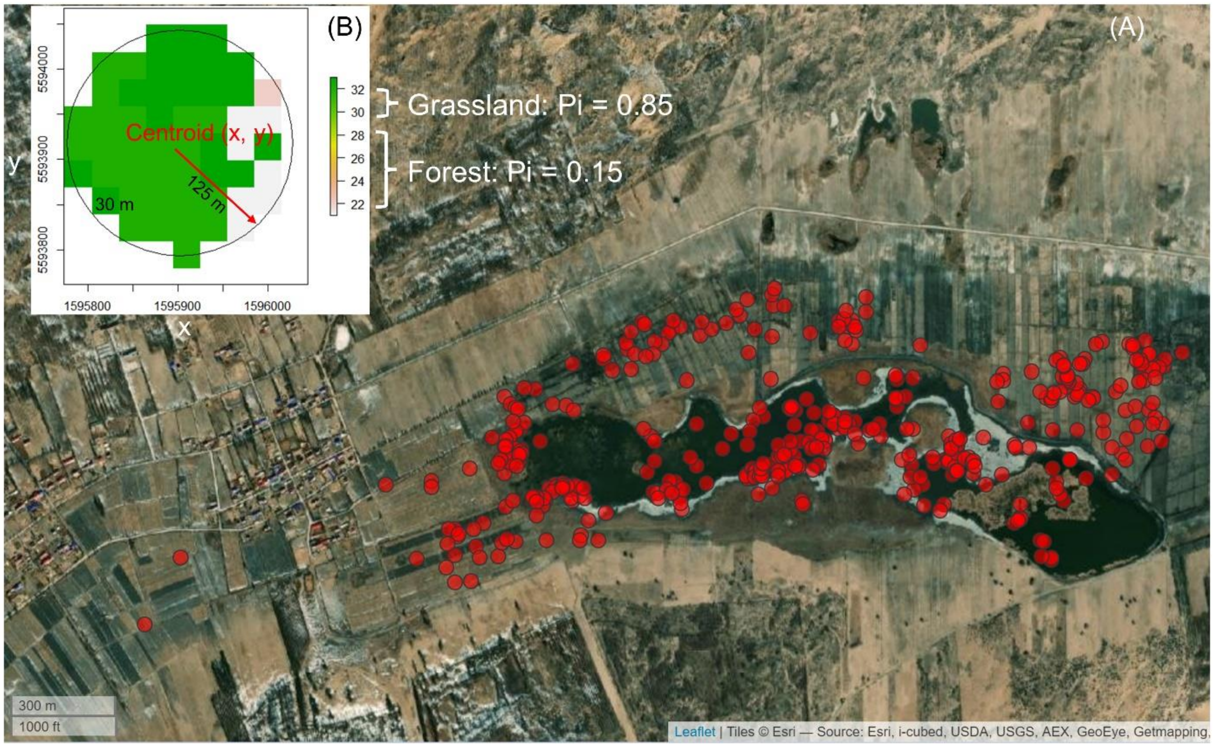


An example of extracting pure pixels in the Northeast China Plain. (A) a pixel shows the mixed-pixel effect (small and fragmented herbaceous areas embedded in large agricultural fields). GPS locations (red dots) represent greater white-fronted geese *Anser albifrons* in the wetlands surrounded by agricultural lands in Khorchin (a major stopover site for waterfowl in the Northeast China Plain). (B) An example of a pure pixel of grassland (Pi = 0.85) in which the value is larger than 0.5.

**Figure S10.**

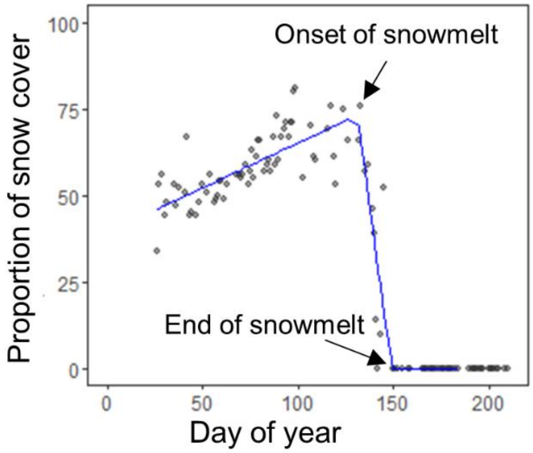


The three-piece segmented regression for a pixel in Lena Delta (the breeding site) for an individual track of greater white-fronted geese *Anser albifrons* in 2019.

**Table S1.**

Summary of sample size, timing of spring migration (in Julian days shown as mean ± SD), and stopover information (mean ± SD) for tracked herbivorous waterfowl ($n$ = 99) in East Asia from 2015–2021.

| Parameters | Arctic-breeding species | | | | Swan goose |
|---|---|---|---|---|---|
| | Tundra swan | Tundra bean goose | Greater white-fronted goose | Lesser white-fronted goose | |
| Track year | 2019-2021 | 2018-2021 | 2015-2021 | 2016-2019 | 2018-2019 |
| Number of individuals | 4 | 28 | 34 | 14 | 19 |
| Number of tracks | 8 | 54 | 63 | 20 | 22 |
| Start date from wintering grounds | 67 ± 9[c] | 67 ± 8[c] | 75 ± 10[b] | 82 ± 7[a] | 71 ± 15[bc] |
| Observed arrival at Northeast China Plain | 87 ± 14[e] | 88 ± 14[d] | 98 ± 12[b] | 100 ± 10[a] | 91 ± 17[c] |
| End date at breeding sites | 149 ± 8[b] | 153 ± 14[b] | 158 ± 13[b] | 172 ± 14[a] | 117 ± 24[c] |
| Migration route | Poyang-Siberia | Poyang-Siberia | Poyang-Siberia | Poyang-Siberia | Poyang-Hulun Lake/Mongolia |
| Migration distance (km) | 5534 ± 267 | 5152 ± 301 | 5242 ± 296 | 5900 ± 640 | 2508 ± 146 |
| Number of stopover sites for each track | 6 ± 3 | 6 ± 1 | 5 ± 2 | 5 ± 2 | 2 ± 1 |
| Length of stay | NCP 56 ± 9<br>Russia 22 ± 12 | NCP 67 ± 21<br>Russia 23 ± 13 | NCP 50 ± 10<br>Russia 21 ± 12 | NCP 46 ± 5<br>Russia 35 ± 12 | NCP 33 ± 22 |

Differences on migration timing for five species were assessed by one way ANOVA followed by Tukey's honestly significant difference (HSD) multiple comparison. Means in a column that do not share a common superscript letter are significantly different ($P < 0.05$).
